# The growth rate of DNA condensate droplets increases with the size of participating subunits

**DOI:** 10.1101/2022.03.29.486311

**Authors:** Siddharth Agarwal, Dino Osmanovic, Melissa A. Klocke, Elisa Franco

## Abstract

Liquid–liquid phase separation (LLPS) is a common phenomenon underlying the formation of dynamic membraneless organelles in biological cells, which are emerging as major players in controlling cellular functions and health. The bottom-up synthesis of biomolecular liquid systems with simple constituents, like nucleic acids and peptides, is useful to understand LLPS in nature as well as to develop programmable means to build new amorphous materials with properties matching or surpassing those observed in natural condensates. In particular, understanding which parameters determine condensate growth kinetics is essential for the synthesis of condensates with the capacity for active, dynamic behaviors. Here we use DNA nanotechnology to study artificial liquid condensates through programmable star-shaped subunits, focusing on the effects of changing subunit size. First, we show that LLPS is achieved in a six-fold range of subunit size. Second, we demonstrate that the rate of growth of condensate droplets scales with subunit size. Our investigation is supported by a general model that describes how coarsening and coalescence are expected to scale with subunit size under ideal assumptions. Beyond suggesting a route toward achieving control of LLPS kinetics via design of subunit size in synthetic liquids, our work suggests that particle size may be a key parameter in biological condensation processes.

## Introduction

The formation of liquid condensates has been recently implicated as a mechanism of control in biological systems ^1^. Liquid liquid phase separation (LLPS) has been observed to involve Chromatin/DNA ^2^, RNA ^3^ and proteins ^4^, and is relevant to a variety of diseases ^5,6^, yet the complexity of natural condensates makes it difficult to harness the mechanisms governing their emergence. This has led to the exploration of platforms for biomimetic condensate design ^7–12^ that have enormous potential toward the development of biological materials that can self-organize, separate, and sort specific components.

A prerequisite to condensate design is the availability of molecular subunits capable of aggregating into distinct dense and dilute phases. While non-specific interactions between subunits are considered to be major drivers in the formation of natural condensates, molecules with programmable specific interactions are of particular interest to engineers and materials scientists. The toolkit of DNA nanotechnology has recently provided a framework to design and build subunits for LLPS. These DNA subunits are star-shaped (nanostars), and assemble from short oligonucleotides that form several double stranded arms. Single stranded domains at the end of the arms, known as sticky-ends, allows subunits to interact ^7,13,14^. Thus, the number of the arms determines the valency of the nanostar, and the length and sequence of the sticky-ends determines the binding affinity among nanostars ^7,15^. Depending on the specific environmental conditions (primarily buffer and temperature), the binding strength varies yielding LLPS or gelation ^14,16^. A major advantage of these DNA nanostars is that sequence design makes it possible to easily customize their structural features, including the sticky end sequence and length, and the number of arms, gaining unprecedented control over the process of LLPS ^7,13–19^. For example, it is possible to build distinct DNA liquid phases and prescribe their interactions ^18^, as well as define their permeability ^17^. These studies point to the possibility of controlling a multitude of macroscopic properties such as condensate size, growth and density, by controlling the nanoscale properties of the phase separating DNA subunits.

Here we present a strategy to gain control over condensate droplet growth rate via the design of DNA subunit size and we demonstrate LLPS via DNA nanostars with arm sizes varying in a 6-fold range. First, we provide a theoretical framework for understanding how subunit size influences the growth rate of droplet condensates. Specifically, we present a ripening model that predicts how coarsening and coalescence scale with subunit size. We then design and experimentally test a series of DNA nanostars differing exclusively in their arm length, which determines the average distance of subunit bonds from their center of mass and thus their apparent diameter ^20^. Via fluorescence microscopy experiments, we observe that condensate growth rate increases as the subunit size increases, we compare empirical observation with the theoretical predictions, and we discuss discrepancies between the two. We hypothesize that the departure from the theory may be due to non-ideal scaling of surface tension and diameter with respect to the nanostar arm size. Our results provide novel nanostar designs that expand the repertoire of DNA motifs for LLPS and will be useful to build complex droplet systems with controllable dynamic behaviors.

## Results

### An ideal theoretical model for droplet growth

We begin by developing a basic mathematical model of droplet growth as a function of time. We consider a system in which molecular subunits of size *σ* interact and separate into two macroscopic distinct phases, a dense phase of concentration *c*_1_ and a dilute phase of concentration *c*_0_ where *c*_1_ > *c*_0_. Phase separation proceeds via the formation of droplets, and we are interested in how the droplet growth kinetics are influenced by subunit size. In the following we shall derive scaling relations for what we term the *ideal model*, where modification of the size of the subunit is matched by commensurate rescaling of *all* the length scales in the system. The two main mechanisms of droplet growth are *coarsening* and *coalescence*. Coarsening refers to the tendency of large droplets to grow at the expense of small ones through constant evaporation and re-condensation of individual particles. Coalescence is the process by which two droplets merge into a single droplet upon coming into contact. The two modes of growth of the radius of the droplets can be described by the Lifshitz-Slyozov scaling law for coarsening ^21^, and through Brownian collision theory for droplet coalescence ^22^. Below we derive a simplified scaling theory for how the growth rates of droplets depends on the size of the subunit *σ*.

The Lifshitz-Slyozov scaling law says growth of the average droplet radius via coarsening can be modeled with the following expression (where we dropped numerical prefactors):

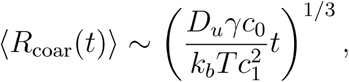

where parameter *k*_*b*_*T* is the Boltzmann constant multiplied by the temperature; *c*_1_ is the concentration of the dense phase; *c*_0_ is the concentration of the dilute phase; *γ* is the surface tension, and *D*_*u*_ is the diffusion constant of the subunit. In the following we shall assume that the Stokes-Einstein relation holds for a single subunit, *D*_*u*_ depends on the subunit diameter: *D*_*u*_ *= k*_*b*_*T* / 3*πησ*.

Brownian collision theory for the growth of droplets via coalescence considers the frequency of interdroplet encounters as a function of droplet size, which depends on how much the droplets are diffusing. The standard expression for the diffusion coefficient *D*_*R*_ of a droplet of radius *R* is given by the Stokes-Einstein relation: *D*_*R*_ *= k*_*b*_*T* / 6*πηR*, with *η* being the viscosity. This is valid for droplets whose thermal motion is driven by collisions of the surrounding solvent with their surface. However, in the case of a droplet comprised of large, permeable subunits (large *σ*), both the interior environment of the droplet and its surface are relatively aqueous. As a result, collisions of the surrounding solvent lead to significant thermal motion within the volume of the droplet as well as motion of its center of mass. The diffusion coefficient of such an object can be approximated by *D*_*R*_ ∼ *D*_*u*_ / *N* where *N* is the number of subunits in the droplet and *D*_*u*_ is the diffusion constant of the subunit ^23^. This means that diffusion should scale as the volume of the droplet. However we note that two droplets of the same size, but constituted of subunits of different size *σ*, will have different *N*. Therefore, given the same droplet size, the diffusion coefficient of these droplets will vary as a function of the subunit size *σ* leading to the scaling *D*_*R*_ ∼ *D*_*u*_*σ*^3^/8*R*^3^, where *σ* is the diameter of the subunit, with *D*_*u*_ *= k*_*b*_*T* / 3*πησ* assuming again the Stokes-Einstein relation holds (the ideal subunit doesn’t have internal thermal motion).

Depending on the size of the subunits, there are two different forms of scaling relationships for coalescence ^24^. For droplets constituted by very small subunits (of the same length scale as the solvent), that are nearly non-permeable, we obtain:

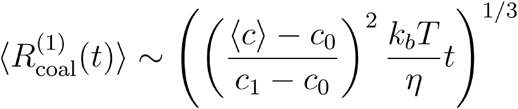

where *η* is the viscosity facing the droplet and ⟨*c*⟩ is the total concentration of material. We deem this *classical Brownian collision theory*. For colloidal droplets constituted by subunits large relative to the solvent, the scaling becomes:

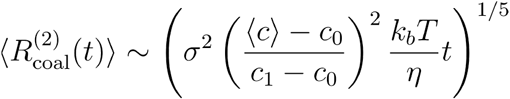

Which we deem *colloidal Brownian collision theory*. These relations show that under the simplest assumptions, droplet growth scales as a power law with respect to time in both coarsening and coalescence-driven growth processes.

To sum up, each mode of growth can be represented as a power law:

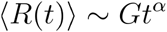

where *G* is a parameter that characterizes how quickly the droplet grows. With the understanding that both coalescence and coarsening may occur simultaneously, in the case of small subunits we expect *α* to be close to ⅓. But for colloidal particles, because it may not be known a priori whether one mode of growth dominates, one should expect *α* to have an arbitrary value between ⅓ and ⅕.

### Theory and computations predict that subunit size influences the growth rate of droplets

We next move to examine the influence of subunit size *σ* on the prefactors of the growth models we just derived. For this purpose, we need to first ask how concentrations in the dense and dilute phases scale with *σ*. To proceed, it is necessary to choose whether to keep a constant *number density* or a constant *volume fraction* in the system as *σ* changes.

We shall first consider the case at fixed volume fraction because it is easier to derive scalings with respect to subunit size assuming a fixed equilibrium point in phase space. Then we shall use those scalings to evaluate growth rates under constant number density, because that is the parameter under experimental control. Specifically, because LLPS equilibrium behavior is typically described as a function of volume fraction *ϕ* = ⟨*c*⟩ (1/6) *πσ*^3^ (rather than the bare number density) and of the effective interaction parameter *ϵ*^∗^ = *k*_*b*_*T* / *ϵ*, where *ϵ* is the interaction energy. This means that holding *ϕ* and *ϵ*^∗^ fixed allows us to control for the observed equilibrium state, while observing changes in growth kinetics due to changes in *σ*.

We can use equilibrium reasoning to derive how we might expect some of these parameters to vary with subunit size *σ*. Imagine two different systems with *σ*_1_ and *σ*_2_ = 2*σ*_1_. In the first case, the equilibrium state will be one large droplet of volume *V*_1_, and a dispersed phase of volume *V*_0_. We imagine taking the equilibrium state of the subunits of *σ*_1_ and doubling every length in the system (it’s important every length gets scaled in the same way). The new system has the same reduced dimensionless parameters as previously, so is at the same point in the phase diagram. In the ideal case, the new system is also at equilibrium, but the volume has increased by a factor of 2^3^ while the total number of particles is held fixed. Then, it must be the case that the concentrations of the dense and dilute phases should scale as *c*_1_ ∼ 1 / *σ*^3^ and *c*_0_ ∼ 1 / *σ*^3^.

Of course, a simpler way of showing the above is asserting that the volume of a subunit itself scales as *σ*^3^, which would yield the same scaling. However this approach is also useful when we next estimate how the surface tension of the droplets scales with *σ*, which is relevant in particular for coarsening.

The canonical form of the surface tension is given by: *γ* = *δF* / *δA* where *F* is the Helmholtz free energy and *A* is the surface area between the dense and dilute phases. While this is in principle a complicated relation, we can proceed again as above, where we know that the final macroscopic state must be consistent through rescaling of all the length scales in the system. In the equilibrium scenario where all length scales in the system are scaled by a factor of *σ* (an affine transformation), all else being unchanged, one derives that *γ*_2_ ∼ *γ*_1_ /*σ*^2^ because the area should increase by the length scale squared, but the macroscopic state remains the same.

By inserting the expressions we just derived into the growth equations, we can draw some conclusions on how condensate droplets are expected to grow as a function of subunit size when keeping a fixed volume fraction:

- Changes in subunit size have no effect on the kinetics of coarsening (with concentration scaling with 1/*σ*^3^, the surface tension *γ* with 1/*σ*^2^, and the subunit diffusion with 1/*σ* all the sigma dependence cancels out)
- For the classical Brownian coalescence theory, rescaling the size of the subunit has no effect on the coalescence kinetics
- For colloidal Brownian coalescence theory, we predict that the growth rate of droplets should scale with the size of the subunit as ∼ *σ*^2/5^

By deriving scalings for fixed number density, we find that:

- Rescaling the size of the subunit has no effect on the kinetics of coarsening
- For Brownian coalescence, the growth rate scales as *σ*^2^
- For colloidal Brownian coalescence, the growth rate scales as *σ*^8/5^

We illustrate the scalings derived above in Fig. 1, where we show computational simulations of colloidal condensate droplet growth for a full particle model in the regime where coalescence dominates (which is where we expect to see a subunit size scaling effect) (see SI section 3). These simulated results confirm that the growth rate of droplets is expected to scale with subunit size *σ*, and the computed scalings agree well with the theoretical scalings derived above.

**Figure 1:**
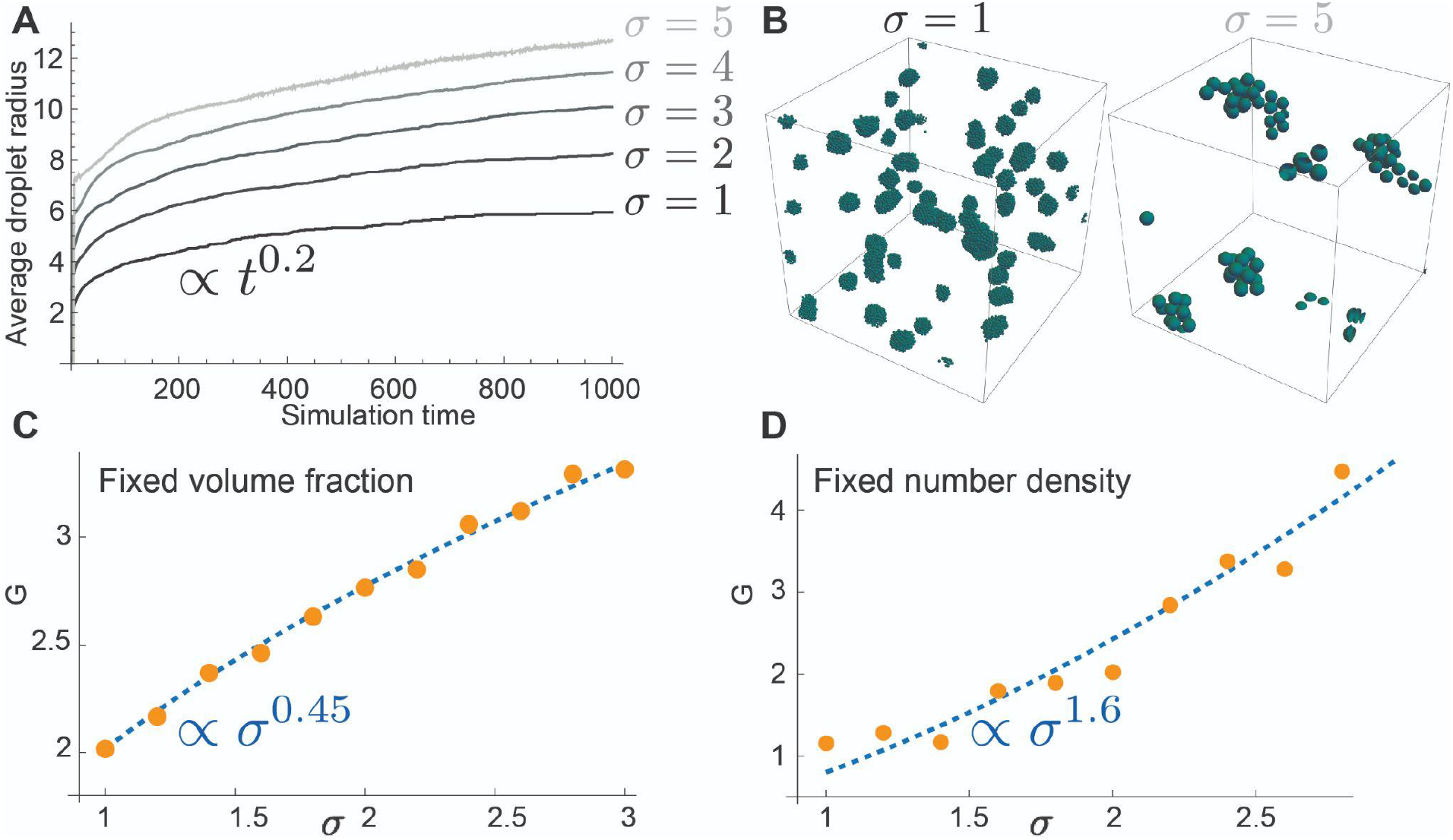
Computational and theoretical exploration of the effect of subunit size on the growth rate of condensates. A) Simulation results showing the average radius of droplet condensates against simulation time for condensates with different subunit size *σ*. The size of the condensate increases faster for larger subunits. B) Simulation snapshots of condensate formation, taken at the end of the simulation in panel A, for subunits with size *σ* = 1 and *σ* = 5. Condensates are comparable in size despite the fact there are fewer particles in the *σ* = 5 simulation. C) values of G calculated from simulation (from ⟨*R*⟩ ∼ *Gt*^*α*^) at a fixed volume fraction against subunit size. The simulation results scale with an exponent of about 0.45, slightly larger than the 0.4 predicted from theory. D) Simulated values of growth rate against subunit size for a fixed number density, this growth rate increases more quickly than for a fixed volume fraction, with an exponent of 1.6, as predicted from the theory.

In the next sections, we describe experiments that investigate how subunit size influences the growth rate of DNA condensate droplets, and later revisit this theory based on experimental observations.

### Design of DNA subunits with programmable size

To test our theoretical and computational predictions, we designed a series of DNA subunits that include three distinct DNA strands assembling into a three arm nanostar motif previously described in the literature ^7,13^. The nanostar subunit includes three double-stranded arms whose junction is made flexible by including unpaired bases (Fig. 2A). These DNA subunits have the capacity to interact with each other via complementary 4-base palindromic ‘sticky end’ (SE) domains present at the end of each arm, which allows distinct nanostars to bind. We tested a collection of subunit designs that differ in their arm length, from 6 bases (∼20.4 nm) to 40 bases (∼136 nm) as depicted in Fig 2C. All these designs share identical SE sequences. To monitor LLPS via fluorescence microscopy, one of the strands was modified to include a fluorescent dye (Cy3) that was mixed at a 10% molar ratio in the solution.

**Figure 2.**
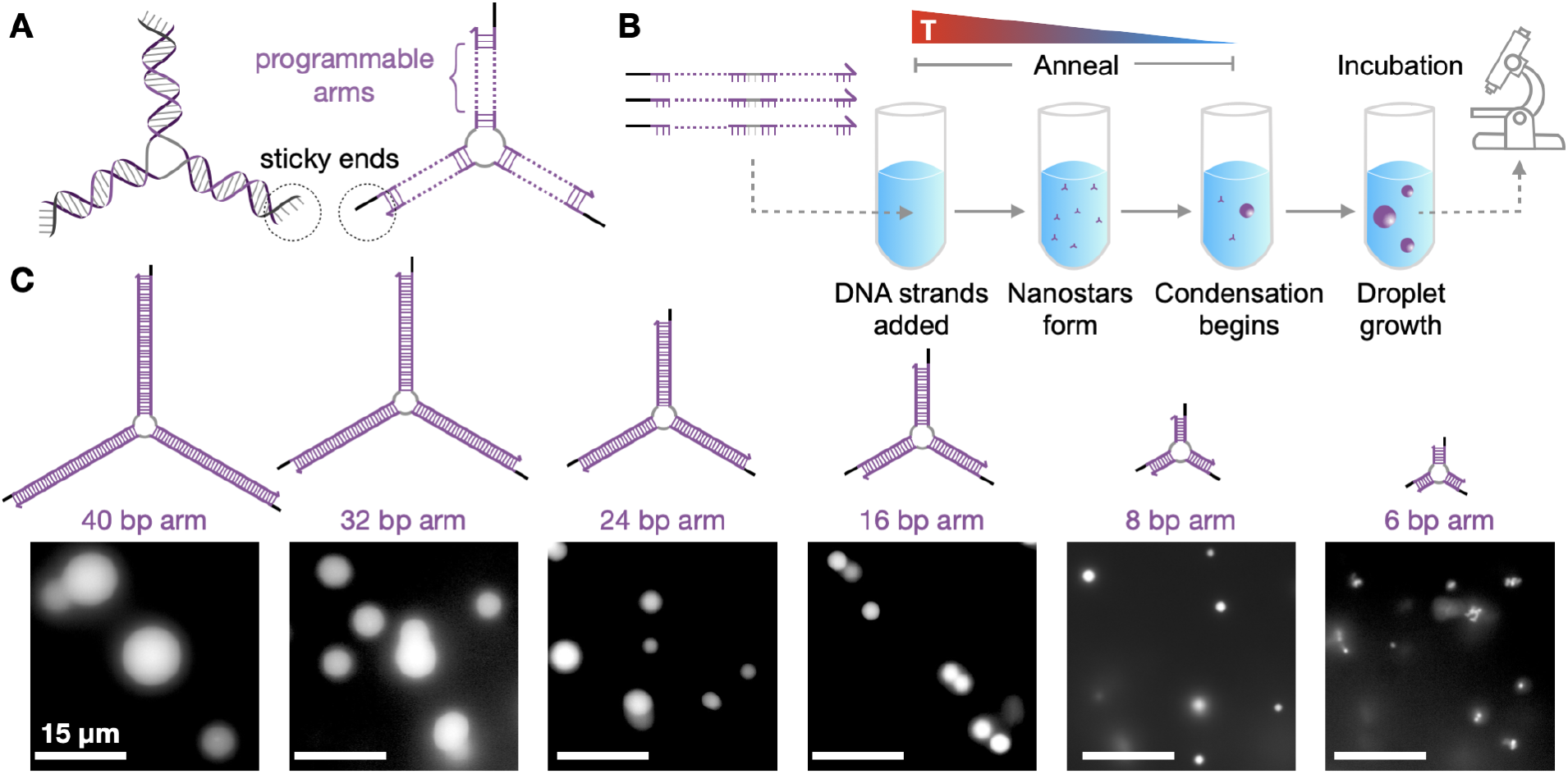
A repertoire of DNA nanostars with distinct arm length yields liquid droplets under consistent experimental conditions. **A:** Schematics of the 3-armed DNA nanostar. We program the length of the nanostar arm while leaving the junction and the sticky end domains unchanged. **B:** Single-stranded DNA oligomers are annealed to form 3-armed DNA nanostructure (nanostars). During the annealing process, nanostars form and at lower temperature interact through their sticky ends and phase separate into condensate droplets. In our experiments, samples are incubated at room temperature and the condensate droplets continue to grow; we take images of the droplets over time via fluorescence microscopy. **C:** We designed nanostars with arm lengths ranging from 40 bases to 6 bases. We show representative fluorescence microscopy images taken right after the annealing step (5 μM nanostar concentration). Scale bars are 15 μm.

We first verified that the DNA subunits undergo LLPS under conditions consistent with previous reports ^7^. Each design was assembled via thermal annealing, from 90° C to 30° C (1°C/min decrease) in 20 mM Tris-HCl (pH 8.0) with 350 mM NaCl. Samples were imaged within one minute of annealing completion, and all designs yielded spherical fluorescent droplets (dense phase) surrounded by dark background (dilute phase). A fraction of droplets showed eccentricity (Fig S7), that is consistent with the presence of fusion events in liquid condensates, as we expected based on the chosen length of the sticky ends and the buffer conditions tested in the literature ^7^. In these experiments it was apparent that the average size of the droplet condensate was proportional to the size of the DNA subunit employed (Fig. 2C). Given that all subunits were annealed with the same protocol, and so with identical incubation times, this initial result already suggests that droplet growth rate depends on subunit size. To quantify the condensate growth rate, we monitored the condensates over time via microscopy and gathered statistics of the temporal changes in condensate droplet size.

### Condensate droplet growth rate depends on subunit size and concentration

After verifying that our DNA nanostar designs have the capacity to yield condensed droplets, we characterized droplet growth at different subunit concentrations. For this purpose, we annealed the DNA subunits at concentrations ranging from 250 µM to 5 µM, and we subsequently allowed the samples to grow at room temperature (set as 27°C in an incubator); aliquots of each sample were taken over a period of 24 hours, and imaged via fluorescence microscopy (Fig 2B). The images were then processed using an automated thresholding process that allowed the extraction of droplet diameter, which was plotted over time (image processing methods are described in SI Section 2). These experiments were focused on nanostars with arm length between 8 and 32 bases, because the small droplets formed by the 6 base design proved challenging to track quantitatively via microscopy (< 1 µm average diameter); in contrast, the droplets generated by the 40 base design coalesced into large condensates occupying most of the field of view before the end of the annealing process (SI Fig. S1).

Fig. 3B shows the average change of droplet condensate diameter during the 60 minutes post anneal, for subunit concentrations of 0.25 µM, 1µM, 2.5 µM and 5 µM. We define the average change in diameter of the condensates as the difference between ‘D_t_’, the average diameter of the condensates measured at time ‘t’, and ‘D_0_’, the average diameter of condensates at the end of the anneal process (considered here our initial condition). For visual comparison, Fig. 3A shows representative images of droplets formed by subunits with different arm length annealed at 1 µM concentration, at different times after annealing is complete (0, 15 and 60 minutes). These data show that, as expected, the average diameter of the condensates increases with the subunit concentration, and confirm the theoretical prediction that subunits with longer arms produce droplets that grow faster. We also measured the number of detected droplets, and found that their total number decreases over time, which is consistent with our expectation of droplet coalescence (SI Fig. S3 and S4). Importantly, we noted a drop in the measured average droplet diameter at longer incubation times: we attribute this to our inability to accurately sample the droplet size distribution due to sedimentation of larger droplets occurring in the incubated sample, which was not mixed or agitated prior to each sampling. For this reason, we narrowed our attention to the droplet growth rate within the first 60 minutes after annealing, when droplet sedimentation should be less prominent as droplets are still relatively small. The change in average diameter over the full 24 hours for different subunit designs at different concentrations is reported in SI Fig. S5 and S6.

**Figure 3.**
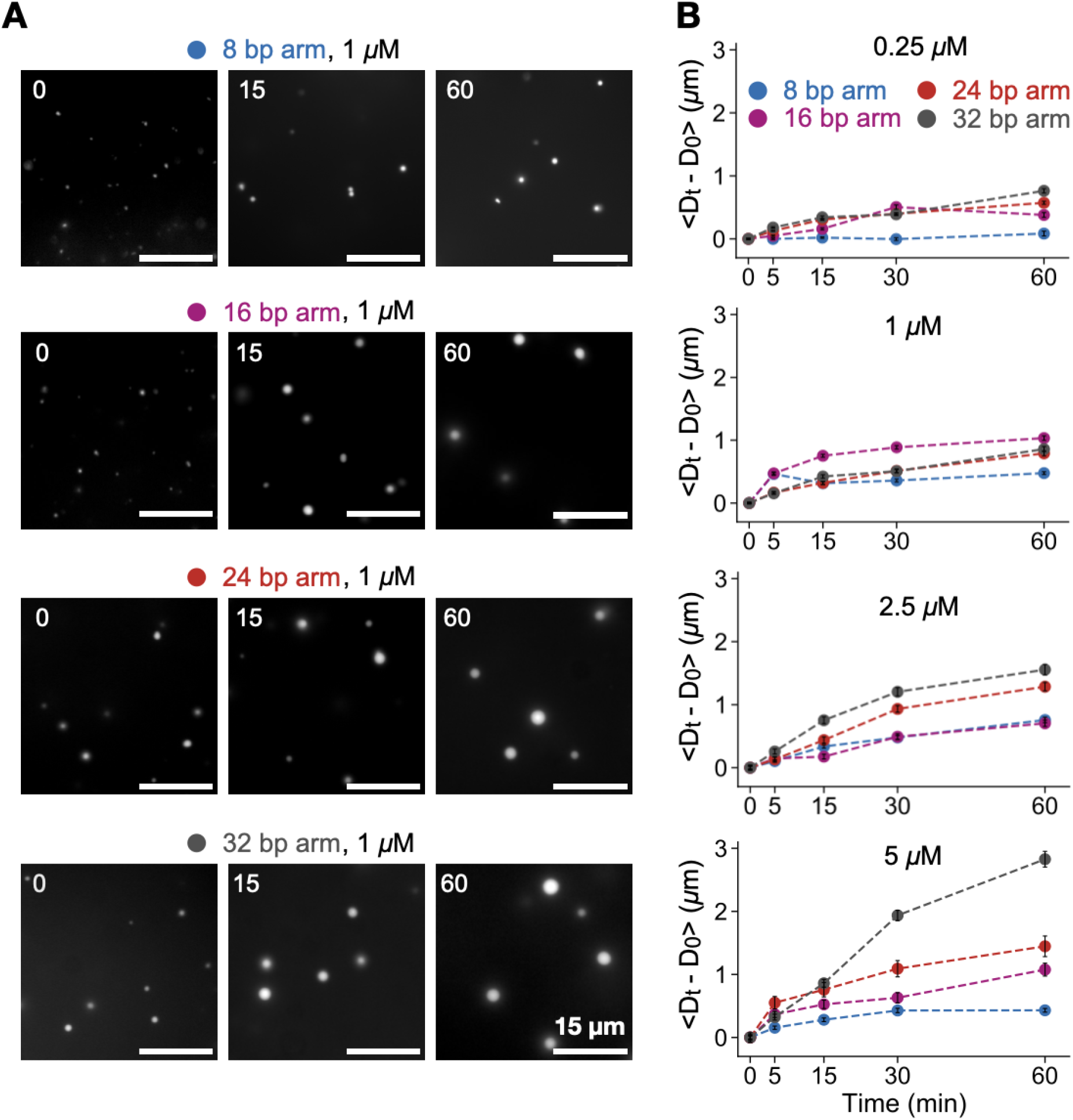
The temporal kinetics of droplet growth are influenced by nanostar arm length and concentration. **A:** Temporal sequence of representative fluorescence microscopy images of condensates formed by nanostar variants having 8, 16, 24 and 32 base arm length, annealed at concentration of 1 µM. The numbers on the top left of the images represent time (in minutes) at which these images were captured from the start of the incubation. Scale bars are 15 μm. **B:** Plots of the increase in diameter of condensates (where D_t_ is the average diameter of the condensates at time ‘t’ and D_0_ is the average diameter of condensates at the end of anneal process) when incubated at room temperature (27° C) over 60 minutes after annealing at subunit concentrations of 0.25 μM, 1 μM, 2.5 μM and 5 μM. The 8, 16, 24, 32 base arm designs are represented in blue, purple, red and black respectively.

### Theoretical and empirical scaling laws of condensate droplet growth

We next seek a more quantitative analysis of how the droplet growth rate scales in our experiments with DNA subunits of different arm length, and compare our analysis to the theory developed earlier. We begin by examining how growth rate scales with time and concentration, for each nanostar design, considering the data in Fig. 3B. We fitted each time trace using the power law *Gt*^*α*^ and determined the effective exponent *α* capturing the change of average droplet size in time (see also SI figure S8); the fitted exponents are reported in Table 1. The average exponent across all the data is *α* ≈ 0.4, which is consistent with a growth model under coarsening or under classical Brownian collision theory.

**Table 1:** Exponents fitted to time traces of average droplet diameter increase (Fig. 3B)

|  |  |  |  |  |
| --- | --- | --- | --- | --- |
|  | 0.25uM | 1.uM | 2.5uM | 5.uM |
| 8 bp arm | 0.34 | 0.24 | 0.46 | 0.31 |
| 16 bp arm | 0.37 | 0.25 | 0.43 | 0.36 |
| 24 bp arm | 0.43 | 0.47 | 0.49 | 0.36 |
| 32 bp arm | 0.45 | 0.50 | 0.61 | 0.50 |

Next, we examine how the average size of droplets scales with subunit size at a single time slice. By fixing our attention on a single time slice, it is easy to compare the measured scaling of growth rate versus subunit size with the ideal scaling under different theories. In this case, we fit the data with a power law of the type *Kσ*^*β*^ and we ask whether data show quantitative consistency with the Brownian particle model or colloidal model for coalescence (the ideal coarsening growth model does not depend on subunit size). We focus our attention on the droplet data collected 60 minutes after anneal (incubation at room temperature), and in Fig. 4 we provide a plot of the average diameter change versus concentration (Fig. 4 A) or nanostar arm length (Fig. 4 C). This particular time point was selected as droplets produced by all subunit variants are sufficiently large for quantitative diameter measurement via epifluorescence microscopy, while we do not expect significant sedimentation and segregation of very large droplets that could bias our measurements.

**Figure 4.**
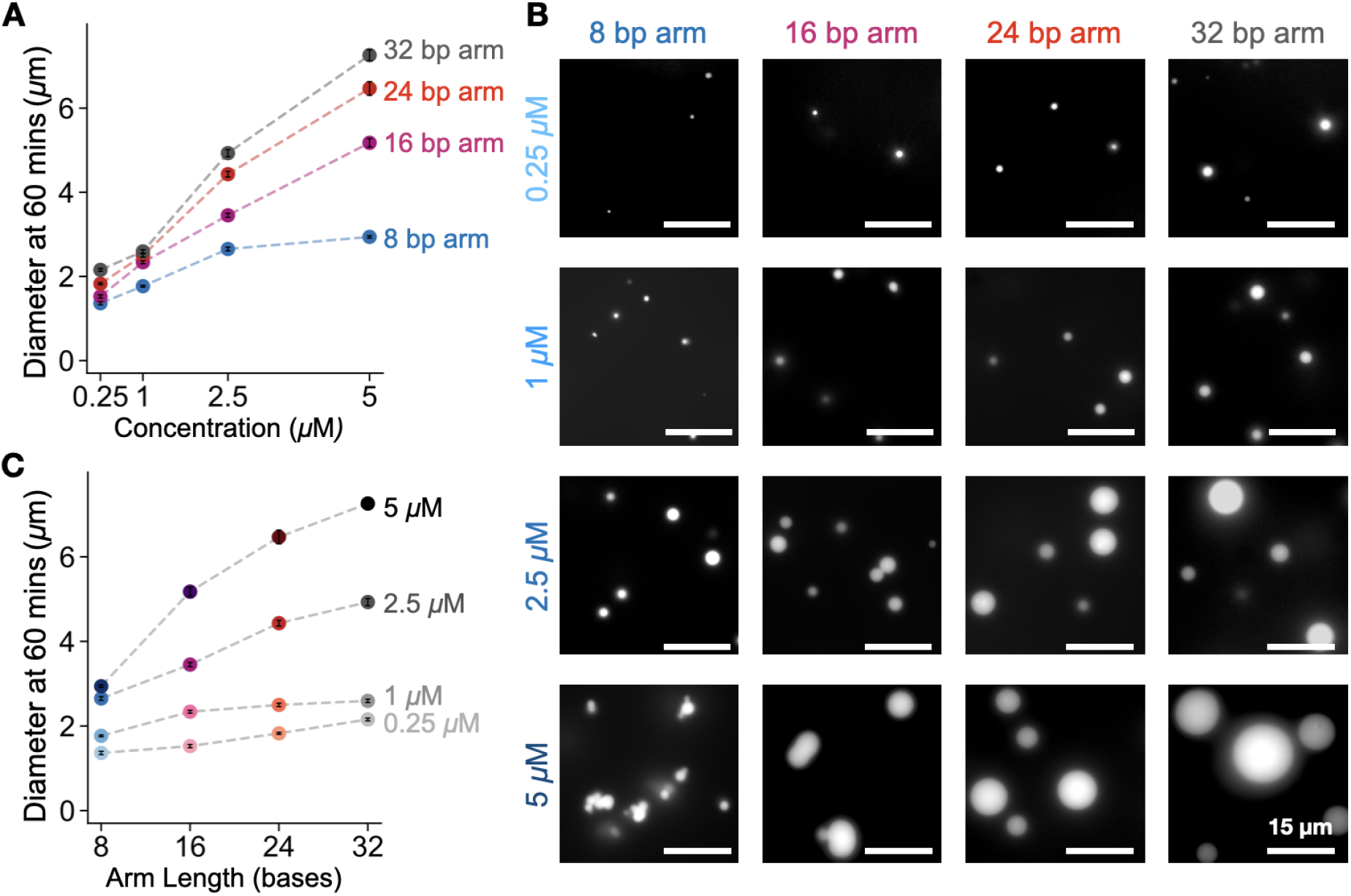
Droplet diameter scales with arm length and concentration. We consider a sample incubated at room temperature (27° C) for 60 minutes after anneal. The 8, 16, 24, 32 base pair arm designs are represented in blue, purple, red and black respectively. **A:** Plots of the average diameter of condensates versus the subunit concentration **B:** Representative fluorescence microscopy images of condensates at time 60mins. Columns represent 8, 16, 24 and 32 arm designs respectively. Rows represent 0.25 μM, 1 μM, 2.5 μM and 5 μM as concentrations respectively. Scale bars are 15 μm. **C:** Plots of the average diameter of condensates versus the arm length. The increase of concentration is marked by a progressively darker shade of each color.

The second column of Table 2 reports the fitted experimental scalings for each concentration. For fixed number density (concentration), the coarsening model does not depend on subunit size, while the Brownian particle model predicts a scaling of *σ*^2^, and the Brownian colloidal model predicts a scaling *σ*^8/5^. The measured scalings point clearly to a dependence of growth from subunit size, but such scalings are significantly lower than the ideal Brownian model scalings, with exception for the 0.25 µM experiments. Yet, we note that the measurements for the 8-base arm length subunits appears qualitatively different to the others (this is particularly evident in log-scale plots in SI Fig. S8); if we ignore the 8-base subunit size, we obtain more consistent fitted exponents (third column of Table 2), that are however still much lower than those predicted by the ideal brownian coalescence theory.

**Table 2:** Scalings of the DNA condensate droplet growth rate observed in Fig. 4C

|  |  |  |
| --- | --- | --- |
| Subunit concentration<br>($\mu\text{M}$) | Experimentally<br>observed scaling against arm<br>length | Experimentally observed scaling<br>excluding 8-arm data |
| 0.25 | 1.78 | 0.34 |
| 1 | 0.36 | 0.10 |
| 2.5 | 0.67 | 0.36 |
| 5 | 0.73 | 0.34 |

These experimental scalings suggest that the ideal theory falls short of describing quantitatively droplet growth in DNA condensates. We formulate some hypotheses that may help explain this discrepancy, by revisiting our theoretical assumptions. First, the coarsening prefactor may depend on subunit size, introducing a dependence of coarsening on subunit size. In the ideal case, we assumed the surface tension *γ* varies with subunit size as 1 /*σ*^2^, and this assumption renders the overall coarsening process independent from subunit size. However, this scaling is justified only if the attractive length varies in precisely the same way as the steric repulsion (as it does in a Lennard-Jones liquid, for example), and in practice this scaling may be more complex. In place of this ideal scaling, one could consider an empirical scaling *γ* ∝ *γ*_0_ / *σ*^2−*β*^, and in this case we would expect the growth rate due to coarsening for different sized subunits to be affected by subunit size as *σ*^*β*^. The fact that the growth rate of droplets increases with subunit size indicates that *β* > 0, meaning that as we increase the size of the DNA nanostar arms, the surface tension doesn’t decrease as much as would be expected from the ideal calculation.

Another possibility is that the size of the subunit, estimated with respect to some control parameter, does not scale precisely as the control parameter. In the case of the DNA nanostar arm, the subunit size was expected to scale as the arm length *L*, which should determine the subunit diameter; however, the arms are short polymers, and for a purely flexible arm this would lead to an effective diameter that would scale as *σ* ∝ *L*^1/2^. Further, we assumed that the nanostar occupies a spherical volume defined by the length of the mobile nanostar arms, yet three-arm nanostars may form a more disc-shaped structure, in which case the scaling of subunit size with respect to polymer arm size would be *σ* ∝ *L*^2/3^. Further investigations would be necessary to test these hypotheses and help refine the theoretical predictions.

Finally, we also note that the growth could possibly be the result of two processes whose relative importance shifts as the size of the subunit changes over time, yielding factors that do not precisely follow one scaling or the other. This possibility is supported by the observed temporal decrease of droplet eccentricity (Fig. S7), which is an indirect measure of the proportion of coalescence events.

## Discussion and conclusion

We have presented theory and experiments showing that the growth rate of DNA condensate droplets is affected by the size of participating subunits. Our theoretical model provides relevant scaling laws under ideal assumptions, and considers coarsening and coalescence, including the colloidal case in which the subunit size is significantly larger than the solvent. Consistently with the literature, the model captures the known fact that droplet growth is driven by a power law with respect to time ^21^; we further demonstrate that a power law also describes the dependence of growth from subunit size. We tested the model predictions by engineering a simple 3-arm DNA nanostar subunit in which the arm-length can be programmably changed, and we verified that larger subunits yield condensate droplets that grow faster over time, confirming qualitatively the trends predicted by the theory. The experimentally observed growth rates however depart from the ideal model, and we offer hypotheses explaining such discrepancy that include a dependence of surface tension on subunit size, as well as a non-linear scaling of subunit diameter relative to the arm-length of the nanostars. Overall, this study delineates a simple strategy to achieve control over the growth rates of DNA-based liquids by simply programming the size of the phase separating subunits, and this approach may be extended to control the kinetics of LLPS of other biomolecules.

Another finding of our work is that all the DNA nanostar designs we tested underwent LLPS without the need for adjustments in the thermal annealing or buffer conditions. While nanostars of different valency have been previously tested ^15^, to our knowledge this is the first systematic demonstration that nanostars with arm length between 6-40 bases have the capacity to yield liquid droplets. Arm length may influence the permeability of the resulting DNA condensate, as previously investigated through the use of linkers that affect the droplet surface tension ^17^. The ability to control droplet permeability is relevant in the context of the potential technological applications of DNA condensates and hydrogels for localization recruitment of other molecules and reactions ^19^. For this purpose, further studies will be needed to characterize the phase diagram as both sticky ends, arm length, and solvent conditions are varied.

While the theory we formulated provides useful guidance on the expected scalings of droplet growth, we found discrepancies between experiments and theoretical predictions of growth rates because the characteristics of DNA subunits likely do not satisfy the ideal assumptions made in our model. Particularly, the surface tension of our DNA droplets may introduce subunit size dependence in the coarsening process; experiments focused on monitoring surface fluctuations for individual droplets may help test this hypothesis. We also believe the subunit diameter may scale non-linearly with the arm length, which is possible given that nanostar arms are held together through a flexible junction. Further, a 3 arm nanostar may not generate an equivalent spherical bond volume, although previous work suggests that 4 arm nanostars can be approximated as such ^20^. Molecular dynamic simulations and scattering experiments could help test this hypothesis experimentally.

Our theoretical analysis suggests that the dependence of condensate droplet growth-rate on subunit size is a general phenomenon. While we are not aware of whether there exist natural biological processes in which condensate growth is regulated by controlling the size of participating molecules, this idea is certainly relevant to the synthesis of artificial condensates and membraneless organelles ^25,26^. By selectively activating or deactivating subunits of different sizes, it may be possible to prescribe how fast a condensate grows depending on context, and it may be possible to program complex systems in which different droplet species grow at distinct rates generating patterns ^18^. Importantly, given a target condensate volume, while using X larger subunits instead of N*X smaller subunits would not change the total mass needed to produce the condensate, using fewer, larger subunits would allow for faster production of the condensate.

We expect that our results will accelerate the design of synthetic biomolecular condensates, in particular custom DNA and RNA liquids, as well as expand the toolkit of composite materials that DNA take advantage of star-shaped subunits ^27,28^, which are relevant for materials science, synthetic biology and biotechnology applications.

## Methods

A detailed description of the methods is included in the SI file that accompanies the manuscript.

### Oligonucleotides

Oligonucleotides were purchased from IDT DNA. Fluorophore-labeled strands were purified to high-performance liquid chromatography (HPLC) grade. All strands more than 60 bases in length were polyacrylamide gel electrophoresis (PAGE) purified. Oligonucleotide sequences and modifications are provided in the SI section ‘Sequences’.

### Sample preparation

All the subunits were formed by mixing the target concentration of each oligomer in a buffer consisting of 20 mM Tris-HCl (pH 8.0) and 350 mM NaCl. One of the strands was modified using a fluorescently labeled dye without sticky ends, which was mixed at a 10% molar ratio in the solution. This solution was annealed using a thermocycler, held at 95°C for 5 min, and then cooled to room temperature at a rate of −1°C/min. After annealing, condensates were incubated at room temperature (set as 27°C in an incubator), unless otherwise specified.

### Fluorescence microscopy

Condensate droplet samples were imaged using an inverted microscope (Nikon Eclipse TI-E) with Nikon CFI Plan Apo Lambda 60X Oil (MRD01605) objective. The Cy3 signal was measured using the Eclipse Cy3 filter cube, using an excitation wavelength of 559 nm.

### Image processing

We extracted DNA condensate size, number, and eccentricity from epifluorescence images using a custom Python script available on Github: https://github.com/klockemel/Condensate-Detection This script requires several Python packages, including scikit-image, pandas, and others ^29, 30, 31^. A complete description of the algorithms used for droplet detection is in SI Section 2.

## Acknowledgements

This research was supported by NSF CAREER award 1938194 to EF and by the Sloan Foundation through award G-2021-16831. We thank Deborah Fygenson for helpful advice and feedback.

