## Supplementary Information for "The growth rate of DNA condensate droplets increases with the size of participating subunits"

### 1. Sequences

The sequences of the oligonucleotide strands were designed starting from the 3 arm nanostars published by (Sato, Sakamoto, and Takinoue 2020) using NUPACK to verify their folding patterns (Zadeh et al. 2011). Standard desalted sequences were ordered from IDT DNA (Coralville, IA, USA). To ensure the efficient formation of motifs, flexibility of the stem was introduced by inserting two spacer bases (TT) at the center of the junction. Bold domains correspond to the subunit sticky ends.

| Name | Sequences (5'-3') |
|---|---|
| Y1_4_6 | **GCG C**CA GTG ATT CTG AGC |
| Y2_4_6 | **GCG C**CC TGT CTT TCA CTG |
| Y3_4_6 | **GCG C**GC TAC GTT GAC AGG |
| Y1_0_6_cy3 | cy3-CAG TGA TTC GTA GC |
| Y1_4_8 | **GCG C**CA GTG AGG TTG TCG TAG C |
| Y2_4_8 | **GCG C**CC TGT CCA TTC CTC ACT G |
| Y2_4_8 | **GCG C**GC TAC GAC TTT GGA CAG G |
| Y1_0_8_cy3 | cy3-CAG TGA GGT TGT CGT AGC |
| Y1_4_16 | **GCG C**CA GTG AGG ACG GAA GTT TGT CGT AGC ATC GCA CC |
| Y2_4_16 | **GCG C**CA ACC ACG CCT GTC CAT TAC TTC CGT CCT CAC TG |
| Y2_4_16 | **GCG C**GG TGC GAT GCT ACG ACT TTG GAC AGG CGT GGT TG |
| Y1_0_16_cy3 | cy3- CAG TGA GGA CGG AAG TTT GTC GTA GCA TCG CAC C |

| | |
|---|---|
| Y1_4_24 | **GCG C**CA GTG AGG ACG GAA GTG AAG GAA CTT GTC GTA GCA TCG CAC CGA CAA AGC |
| Y2_4_24 | **GCG C**GT CGC ATC CAA CCA CGC CTG TCC ATT GTT CCT TCA CTT CCG TCC TCA CTG |
| Y2_4_24 | **GCG C**GC TTT GTC GGT GCG ATG CTA CGA CTT TGG ACA GGC GTG GTT GGA TGC GAC |
| Y1_0_24_cy3 | cy3- CAG TGA GGA CGG AAG TGA AGG AAC TTG TCG TAG CAT CGC ACC GAC AAA GC |
| Y1_4_32 | **GCG C**CA GTG AGG ACG GAA GTG AAG GAA CTC TCC GCG TTG TCG TAG CAT CGC ACC GAC AAA GCG AAC ACG T |
| Y2_4_32 | **GCG C**GC CTC TGT GTC GCA TCC AAC CAC GCC TGT CCA TTC GCG GAG AGT TCC TTC ACT TCC GTC CTC ACT G |
| Y2_4_32 | **GCG C**AC GTG TTC GCT TTG TCG GTG CGA TGC TAC GAC TTT GGA CAG GCG TGG TTG GAT GCG ACA CAG AGG C |
| Y1_0_32_cy3 | cy3- CAG TGA GGA CGG AAG TGA AGG AAC TCT CCG CGT TGT CGT AGC ATC GCA CCG ACA AAG CGA ACA CGT |
| Y1_1_40 | **GCG C**CA GTG AGG ACG GAA GTG AAG GAA CTC TCC GCG TCT CCG CGT TGT CGT AGC GTC GTA GCA TCG CAC CGA CAA AGC GAA CAC GT |
| Y1_2_40 | **GCG C**GC CTC TGT GTC GCA TCC AAC CAC GCC TGT CCA CCT GTC CAT TCG CGG AGA CGC GGA GAG TTC CTT CAC TTC CGT CCT CAC TG |
| Y1_3_40 | **GCG C**AC GTG TTC GCT TTG TCG GTG CGA TGC TAC GAC GCT ACG ACT TTG GAC AGG TGG ACA GGC GTG GTT GGA TGC GAC ACA GAG GC |
| Y1_0_40_cy3 | cy3- CAG TGA GGA CGG AAG TGA AGG AAC TCT CCG CGT CTC CGC GTT GTC GTA GCG TCG TAG CAT CGC ACC GAC AAA GCG AAC ACG T |

### 2. Methods

**Oligonucleotide preparation**

Oligonucleotides were purchased from IDT DNA. Fluorophore-labeled strands were purified to high-performance liquid chromatography (HPLC) grade. All strands more than 60 bases in length were polyacrylamide gel electrophoresis (PAGE) purified. Oligonucleotide sequences and modifications are provided in the SI section ‘Sequences’.

**Assembly of motifs using thermal annealing:**
All the motifs were formed by mixing the desired concentration of each component oligomer in a buffer consisting of 20 mM Tris-HCl (pH 8.0) and 350 mM NaCl. To fluorescently label the condensates, one of the strands was modified using a fluorescently labeled dye without SEs, which was mixed at a 10% molar ratio in the solution. This solution was placed in a thermocycler, held at 95°C for 5 min, and then cooled to room temperature at a rate of −1°C/min. Once the anneal process was over, condensates were allowed to grow at room temperature (set as 27°C in an incubator), unless otherwise specified.

**Preparation of the observation chamber:**
Coverslips (Fisherbrand™, cat: 12-545-JP) measuring 60 x 22 mm, with a thickness between 0.13 to 0.17 mm were soaked in 5% (w/v) bovine serum albumin (BSA) and dissolved in 20 mM Tris-HCl (pH 8.0) for over 30 min to prevent nonspecific interactions of the DNA on the glass surface. After the BSA coating, the glass slides were washed two times with distilled water and dried under an airflow. A square parafilm (Parafilm M® from Fisher Scientific, cat:S37440) slice with a punched hole in the middle was stuck to the BSA-coated glasses by heating the slides to 50°C for 1 min before imaging. After the coverslips returned to room temperature, 2.5uL of sample was pipetted out in the punched hole. Another smaller coverslip (Fisherbrand™, cat: 12-545-AP) measuring 30 x 22 mm, with a thickness between 0.13 to 0.17 mm was placed on top of the parafilm to avoid evaporation of the sample solution during the observation period. The BSA coating causes partial dewetting which slows down the process of droplets adhesion to the surface, and allows us to observe near-spherical DNA droplets.

**Fluorescence microscopy**
Droplet samples were imaged using an inverted microscope (Nikon Eclipse TI-E) with Nikon CFI Plan Apo Lambda 60X Oil (MRD01605) objective. The Cy3 signal was measured using the Eclipse Cy3 filter cube, using an excitation wavelength of 559 nm.

**Image processing**

We extracted DNA condensate size, number, and eccentricity measurements from epifluorescence micrographs using a custom Python script available on Github:

https://github.com/klockemel/Condensate-Detection

This script implements several Python packages, including scikit-image, pandas, and others (van der Walt et al. 2014, Reback et al. 2021, McKinney 2010).

Epifluorescence images capture objects both inside and outside of the plane of focus. Objects outside of the plane of focus may be a different size or shape than they appear in an image and have softer edges than objects in the plane of focus. To maintain confidence in our measurements of condensate characteristics, we sought to measure only objects within the plane of focus by implementing an edge-based threshold method. First the image is smoothed with a Gaussian filter to limit the influence of noise inherent to a fluorescence micrograph in detection of condensates. A Sobel filter is then applied to the smoothed image to find the edges within the image. An Otsu threshold is used to separate condensate edges from the background of the image, followed by binary operations to clean the resulting binary image of edges. The image is thinned such that each feature or edge is 1-pixel thick. Enclosed edges are then filled with a binary fill holes method. A binary opening of the image removes any unenclosed regions such as lines or speckles. Finally, the image is dilated using a disk of radius 5 pixels. Without the dilation, objects larger than about 1.5 um in diameter are systematically underestimated in the threshold process. As a majority of the condensates observed in this work are larger than 1.5 um in diameter, we chose to include the dilation although it systematically overestimates small objects. A watershed algorithm is applied to the cleaned image to separate condensates which may be near to or touching other condensates. Finally, the area, diameter, and eccentricity for individual condensates are measured, as well as the total number of condensates. Diameter is estimated as the diameter of a circle with the same area as a given condensate. All user-input parameters for each image are saved in a csv file, and a diagnostic image with labeled condensates is generated. We processed 8 images (identical size of field of view) for each time point and condition. Within a set of 8 images, if there were less than 16 condensates with diameter 1.5 um or smaller each, or two condensates of that size per image, we considered the number of condensates for that condition to be 0 and did not plot the data. For the supplementary data on condensate eccentricity, the measurements were extracted using the same method described above without watershed segmentation to separate nearby or touching condensates, as touching or joining condensates are what we sought to measure with this characteristic.

### 3. Computational simulations

In order to lend credence to the theoretical scalings derived in the main paper, we perform coarse grained simulations on 10000 Lennard Jones potential interacting particles. These particles interact via the following potential:

$$\phi(r) = 4\epsilon\left(\left(\frac{\sigma}{r}\right)^{12} - \left(\frac{\sigma}{r}\right)^{6}\right),$$

where $\epsilon$ is the strength of attraction between particles and $\sigma$ is their diameter. $\epsilon$ is the strength of attraction between particles and $\sigma$ is their diameter.

We perform these simulations with a Langevin thermostat in an NVT ensemble. The results of the simulations are summarized in Fig. 1 of the main text. In order to observe the effect of coalescence dynamics (where we predict that there should be an impact due to subunit size) these simulations are performed in the regime where coalescence should dominate over coarsening, which we do by setting the interaction strength of $\epsilon = 2.5 k_b T$.

### 4. Additional data and experiments

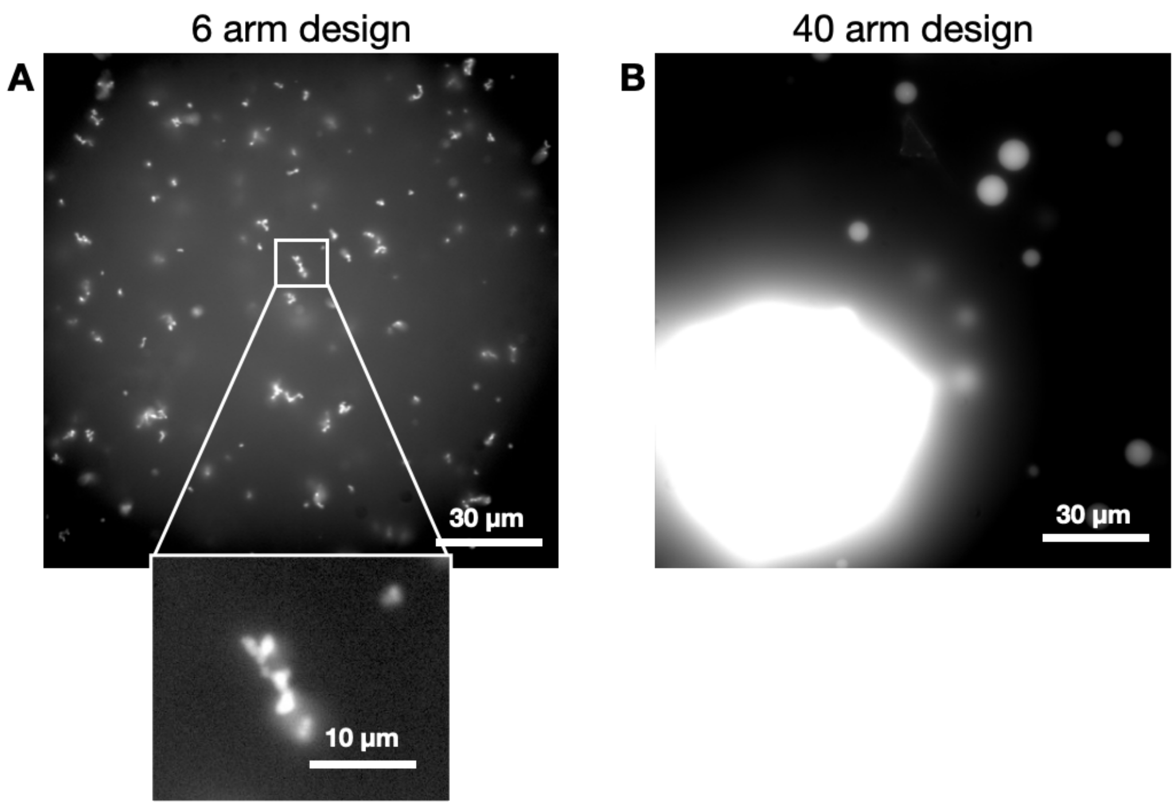


**Figure S1: Microscopy images representing the DNA condensates formed after anneal** (A) 6 base-pair length arm and (B) 40 base-pair length arm design. Inset image in A shows a magnified version of the 6 base-pair length arm design condensates. Non-spherical droplets are the result of slow coalescence events; plots of droplet eccentricity are in Fig. S7. Scale bars as represented.

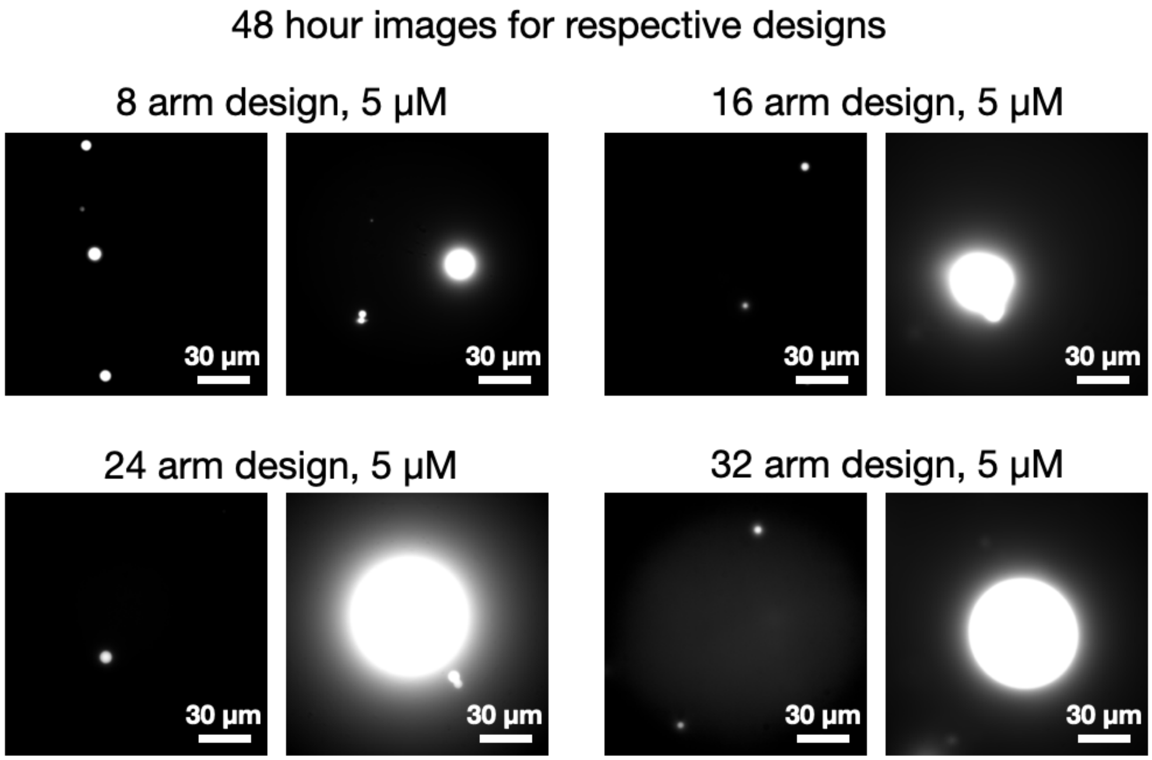


**Figure S2: Microscopy images representing the DNA condensates formed after anneal and 48 hours after incubation at room temperature (27° C).** Scale bars are 30 μm.

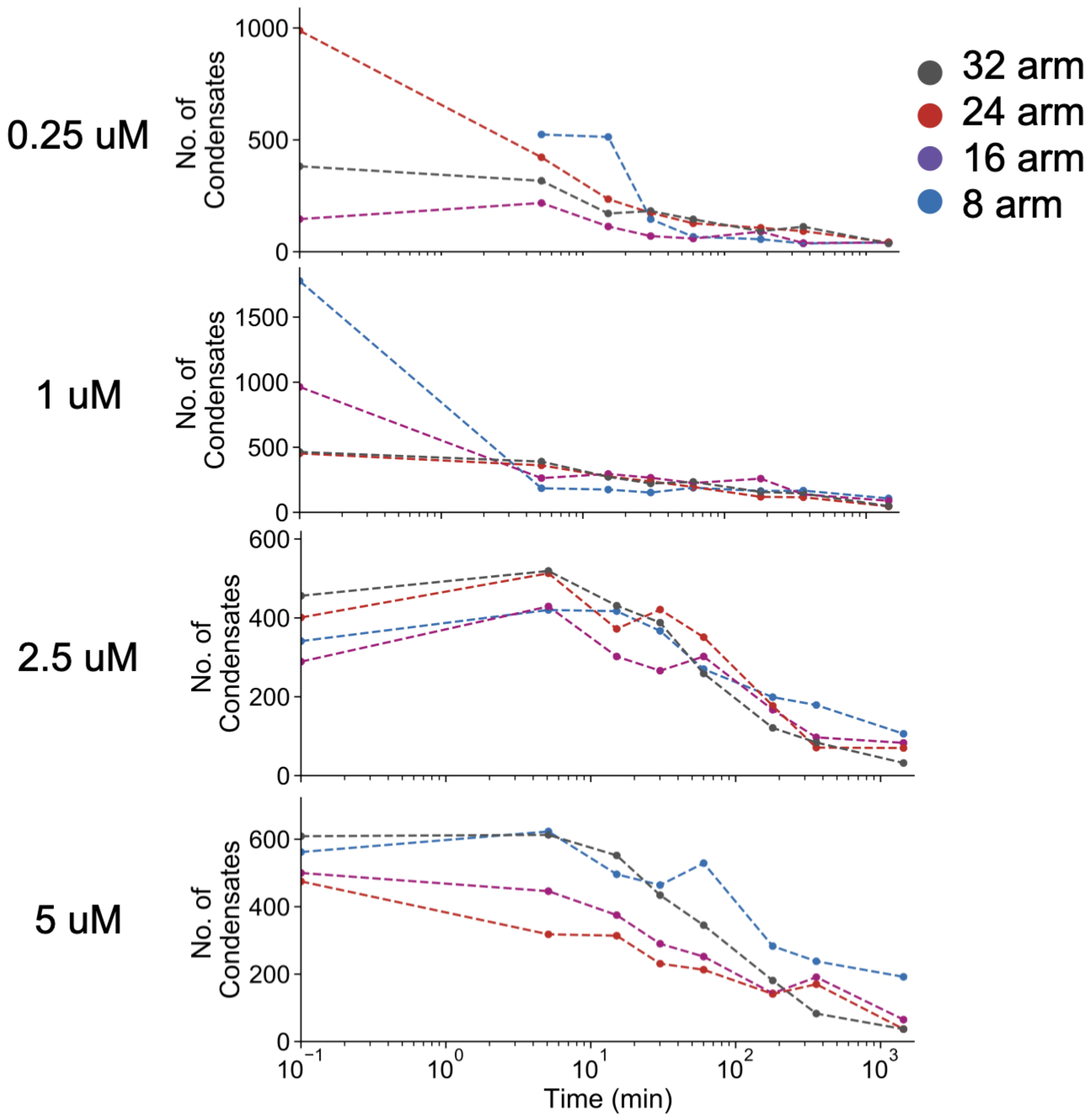


**Figure S3: Number of droplets detected, organized by nanostar concentration**. These are the condensate droplets detected and used to compute the average condensate diameter in each experiment. Data are organized by monomer/subunit concentrations of 250 nM, 1000 nM, 2500 nM and 5000 nM. The 8, 16, 24, 32 base pair arm designs are represented in blue, purple, red and black respectively with increasing values of concentration denoted in progressively darker shades of the color.

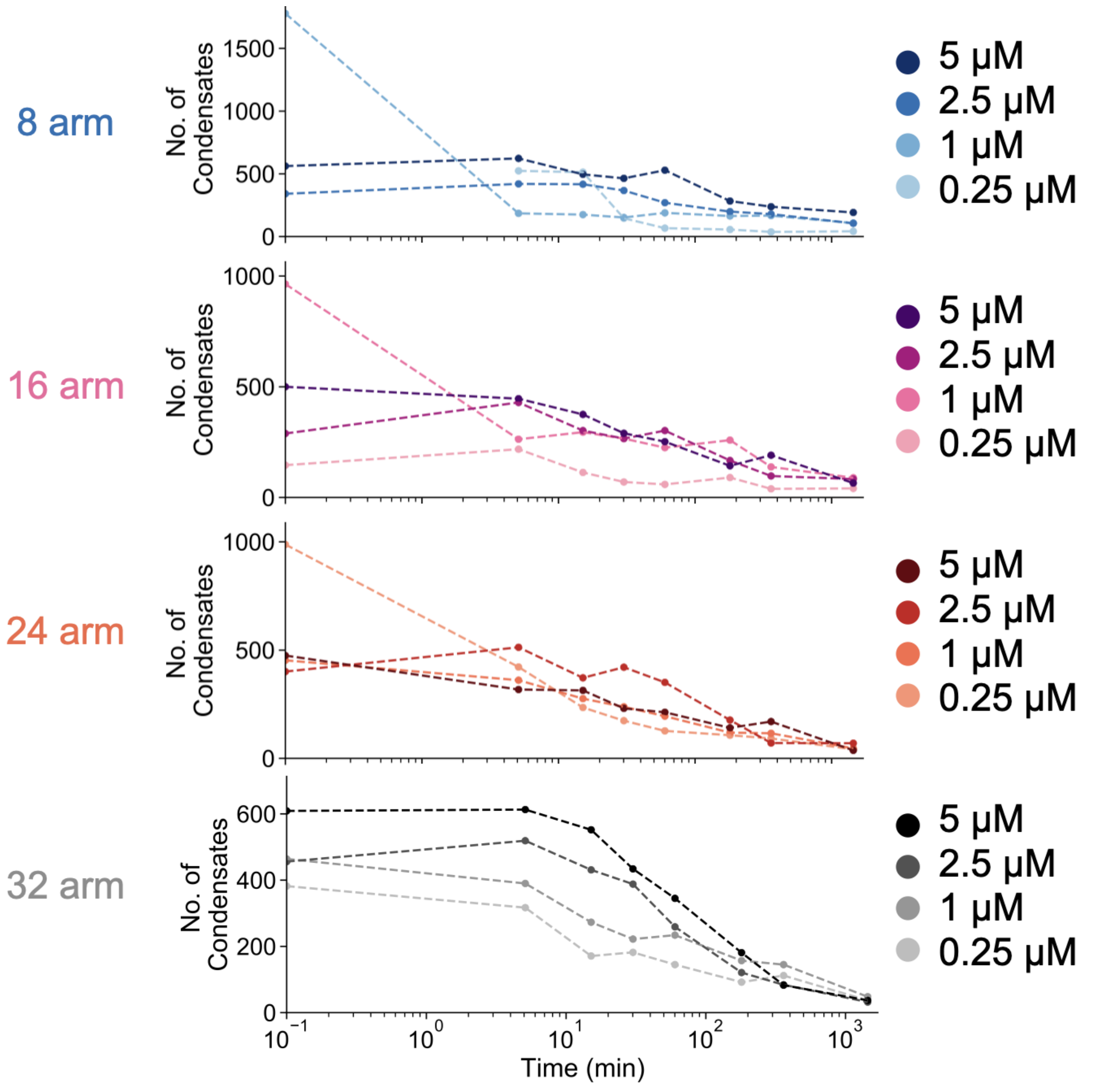


**Figure S4: Number of droplets detected, organized by nanostar arm-length**. These droplets were used to compute the average diameter of the condensates in each experiment. This is the same data shown in Fig. S3, but organized by nanostar arm length. The 8, 16, 24, 32 base pair arm designs are represented in blue, purple, red and black respectively with increasing values of concentration denoted in progressively darker shades of the color.

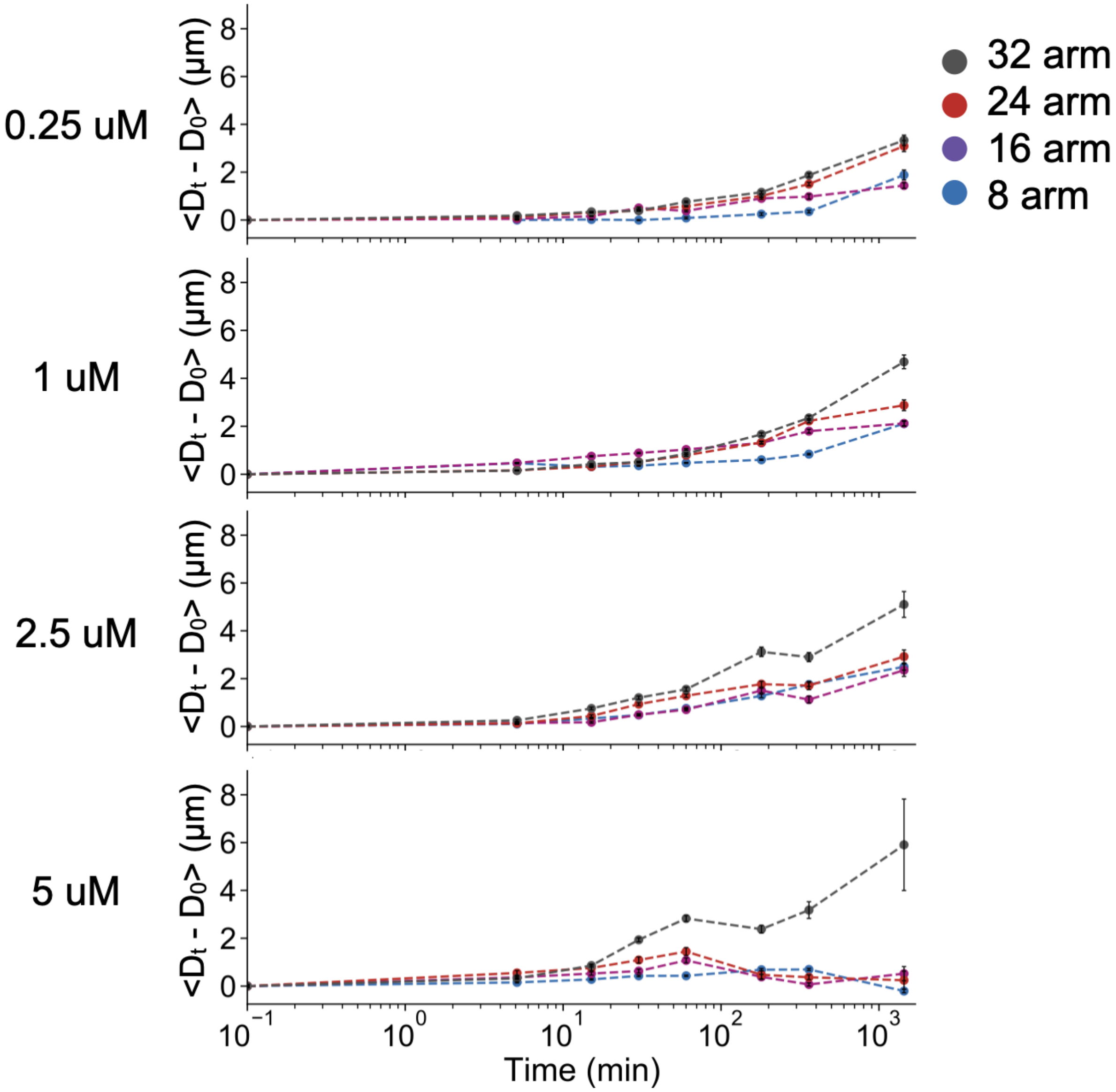


**Figure S5: Increase in diameter of condensates over 24 hours, organized by nanostar concentration.** Here $D_t$ is the average diameter of the condensates at time 't' and $D_0$ is the average diameter of condensates at the end of anneal process); samples when incubated at room temperature (27° C) over 24 hours. Data is organized by monomer/subunit concentrations of 250 nM, 1000 nM, 2500 nM and 5000 nM. The 8, 16, 24, 32 base pair arm designs are represented in blue, purple, red and black respectively.

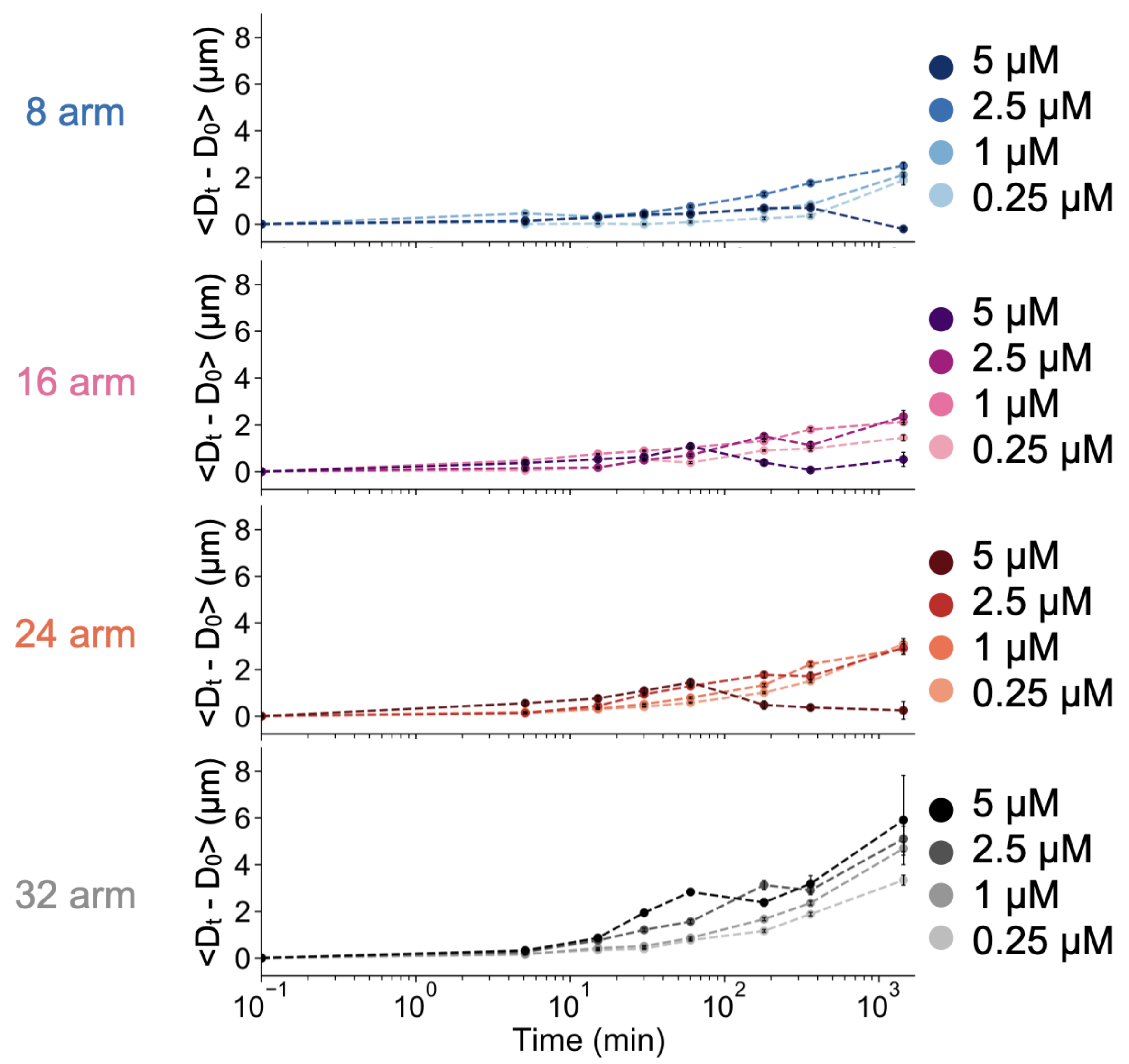


**Figure S6: Increase in diameter of condensates over 24 hours, organized by nanostar arm-length.** $D_t$ is the average diameter of the condensates at time 't' and $D_0$ is the average diameter of condensates at the end of anneal process; samples were incubated at room temperature (27° C) over 24 hours. The 8, 16, 24, 32 base pair arm designs are represented in blue, purple, red and black respectively with increasing values of concentration denoted in progressively darker shades of the color.

#### Eccentricity of droplets

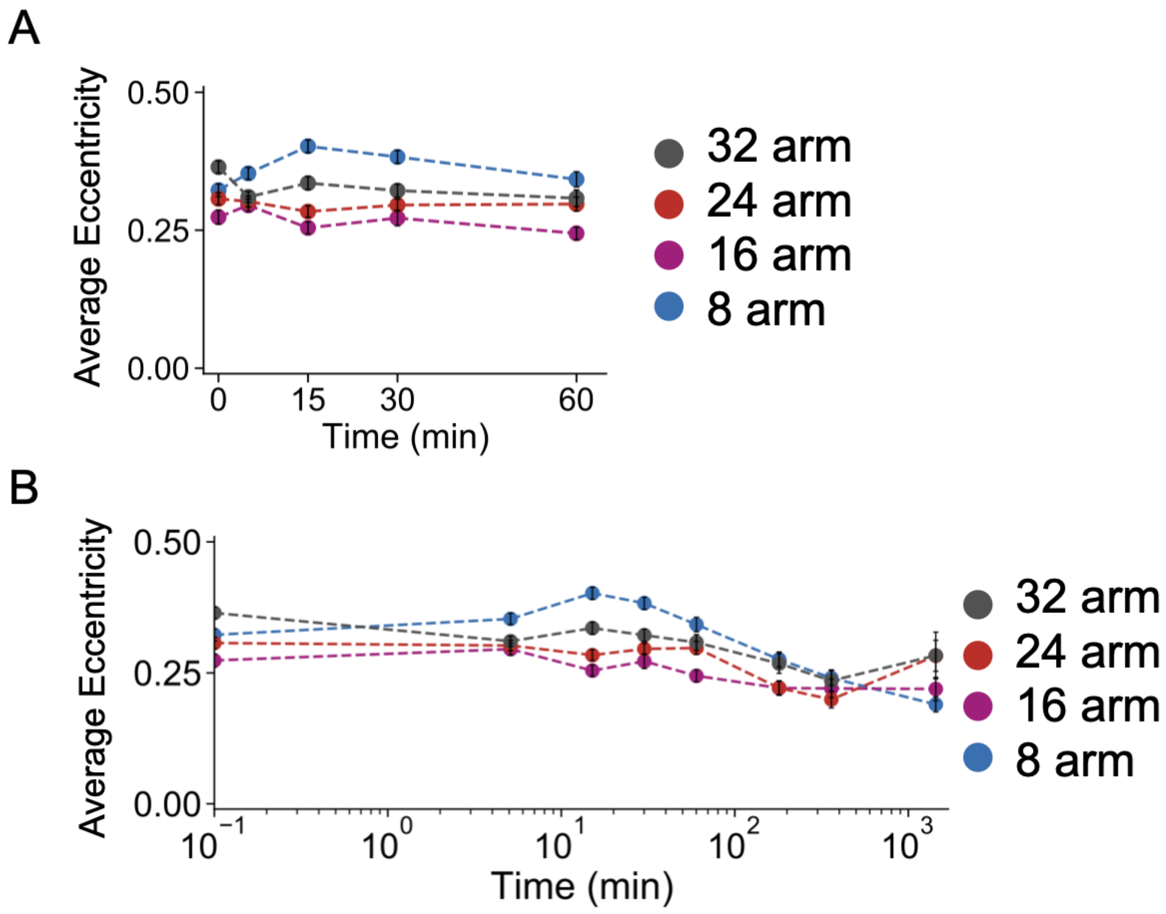


**Figure S7: Average eccentricity of condensates for the 2.5 uM experiment** through 60 min (A) and 24 hours (B) when incubated at room temperature (27° C). The 8, 16, 24, 32 base pair arm designs are represented in blue, purple, red and black respectively.

The eccentricity of droplets gives us information about two characteristics of our system: 1) coalescence events and 2) the time it takes for fusion events to be completed (relaxation time scale associated with fusion). Larger eccentricity indicates that the system is growing more through coalescence, or that it takes longer for droplets to fuse and return to a spherical shape. The data in Fig. S7 indicates that eccentricity is initially largest for 8 arm nanostars; later on in the experiment eccentricity increases as a function of arm length for all other nanostars. The increase in eccentricity for arms 16, 24, and 32 is consistent with the fact that a longer arm length leads to larger volume fraction, thus making encounters between droplets more likely. The anomalous 8 arm data suggests that relaxation times for fusing droplets is larger than expected when there is a large number of small droplets; eccentricity decreases later on as larger droplets have formed but the smaller volume fraction for 8 arm nanostars should lead to less droplet encounters.

#### Fitting Routines for Time Series Data

In order to determine the exponents of time series data we apply a fitting routine to the experimental data in log-log space. As fitting is rather sensitive to variations induced by single points for the 8 time points we performed measurements, we develop a routine to exclude anomalous readings and extract the exponent in the following way:

- The data is transformed to a log-log scale
- On a log log scale, power laws should appear linear, we employ a least squares linear fitting routine to the data on the log-log scale
- If a point included in the data leads to large variations in the fitted exponent (calculated by looking at variance of the residuals norm) it is excluded from the fitting routine. If the points are all fit perfectly by a straight line, no points are excluded
- The slope of the line is the exponent

The fitting was done in Mathematica, and the results are shown in Fig. S8.

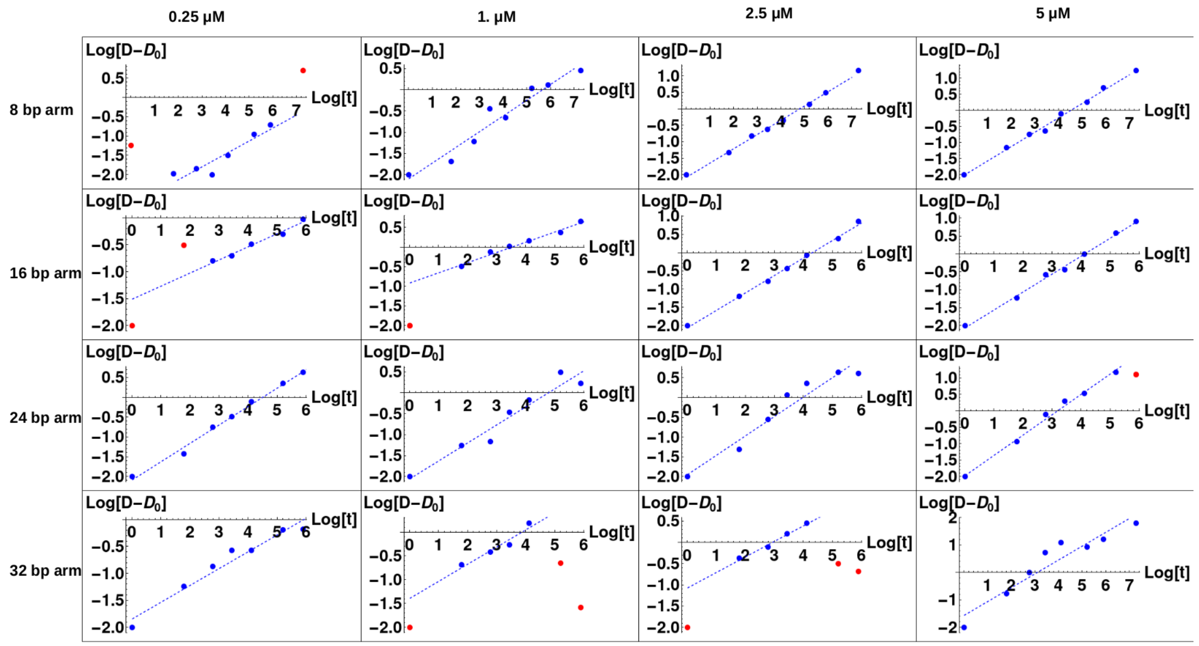


Figure S8: Results of fitting routines for time series data; the blue dots are experimental measurements, the dashed line is the line of best fit, and the points that the algorithm excludes are colored in red.

Walt, Stéfan van der, Johannes L. Schönberger, Juan Nunez-Iglesias, François Boulogne, Joshua D. Warner, Neil Yager, Emmanuelle Gouillart, Tony Yu, and scikit-image contributors. 2014. “Scikit-Image: Image Processing in Python.” *PeerJ* 2 (June): e453.

Zadeh, Joseph N., Conrad D. Steenberg, Justin S. Bois, Brian R. Wolfe, Marshall B. Pierce, Asif R. Khan, Robert M. Dirks, and Niles A. Pierce. 2011. “NUPACK: Analysis and Design of Nucleic Acid Systems.” *Journal of Computational Chemistry* 32 (1): 170–73.
